# Dynamic inhibition of sensory responses mediated by an olfactory corticofugal system

**DOI:** 10.1101/2020.03.29.014571

**Authors:** Renata Medinaceli Quintela, Jennifer Bauer, Lutz Wallhorn, Daniela Brunert, Markus Rothermel

## Abstract

Processing of sensory information is substantially modulated by centrifugal projections from higher cortical areas, yet their behavioral relevance and underlying neural mechanisms remain unclear in most cases. The anterior olfactory nucleus (AON) is part of the olfactory cortex and its extensive connections to lower and higher brain centers put it in a prime position to modulate early sensory information in the olfactory system. Here, we show that optogenetic activation of AON neurons in awake animals was not perceived as an odorant equivalent cue. However, AON activation during odorant presentation reliably suppressed odor responses. This AON mediated effect was fast and constant across odors and concentrations. Likewise, activation of glutamatergic AON projections to the olfactory bulb (OB) transiently inhibited the excitability of mitral/tufted cells (MTCs) that relay olfactory input to cortex. Single-unit MTC recordings revealed that optogenetic activation of glutamatergic AON terminals in the OB transiently decreased sensory-evoked MTC spiking, regardless of the strength or polarity of the sensory response. These findings suggest that glutamatergic AON projections to the OB suppress early olfactory processing by inhibiting OB output neurons and that the AON can dynamically gate sensory throughput to the cortex.

**Significance Statement:** The anterior olfactory nucleus (AON) as an olfactory information processing area sends extensive projections to lower and higher brain centers but the behavioral consequences of its activation have been scarcely investigated. Using behavioral tests in combination with optogenetic manipulation we show that in contrast to what has been suggested previously, the AON does not seem to form odor percepts but instead suppresses odor responses across odorants and concentrations. Furthermore, this study shows that glutamatergic cortical projections to the olfactory bulb suppress olfactory processing by inhibiting output neurons, pointing to a potential mechanisms by which the olfactory cortex can actively and dynamically gate sensory throughput to higher brain centers.

**Highlights:** AON stimulation suppresses odor responses across odorants and concentrations

AON activation is not perceived as an odorant equivalent cue

The AON dynamically shapes olfactory bulb output on a fast timescale

AON input to the olfactory bulb strongly suppresses mitral/tufted cells firing

## Introduction

The ability to perceive external information via sensory systems is crucial for an animal in order to navigate and survive in a complex environment. In a classical view, the brain processes sensory information solely based on a hierarchical organization where sensory information is shaped and refined by subsequent processing steps. However, in order to guarantee appropriate, flexible and fast reactions in a rapidly changing environment, it is beneficial to implement additional mechanisms that modulate information in a situation dependent fashion. One way to do so are cortical top-down projections, where sensory information is received from sensory cortices, and processed information is then transmitted to lower sensory processing centers to modulate incoming sensory signals. Understanding the neural mechanisms underlying sensory perception thus requires understanding the neural circuits involved in both bottom-up and top-down mechanisms.

One prominent center of cortical top-down projections in olfaction is the anterior olfactory nucleus (AON), an olfactory cortical area located in the forebrain just caudally of the olfactory bulb (OB), the first relay station of olfactory signals within the brain. The AON can be divided into two distinct zones, pars externa, a thin ring of cells in the rostral part of the AON, and pars principalis, containing the majority of AON cells (Valverde et al., 1989; Brunjes et al., 2005). Its extensive connectivity with primary and secondary processing centers (see (Brunjes et al., 2005)) and its position as both a “bottom-up” relay of ascending sensory input from the OB and a source of “top-down” input to the OB render the AON an interesting model system for investigating higher-order olfactory processing and the interplay of ascending and descending information.

The AON is the largest source of cortical projections to the OB (Carson, 1984; Shipley and Adamek, 1984). AON-derived axons have been shown to project to multiple layers of the OB (Reyher et al., 1988; Padmanabhan et al., 2016; Wen et al., 2019). This includes the granule cell layer which contains the majority of inhibitory interneurons of the OB, as well as the layers containing the output neurons of the OB, the external plexiform and the mitral cells layer. Furthermore, AON projections are bilateral, i.e. the AON does not only send axons to the ipsilateral but also, via the anterior commissure, to the contralateral OB (Brunjes et al., 2005; Illig and Eudy, 2009). Similar to cortical back projections from piriform cortex (Boyd et al., 2015; Otazu et al., 2015), the AON was shown to send sensory evoked feedback to the OB (Rothermel and Wachowiak, 2014).

The AON has been implicated in a range of different functions, including serving as the first site of integrated odor percept formation (Haberly, 2001; Wilson and Sullivan, 2011), olfactory memory (Haberly, 2001; Aqrabawi and Kim, 2018a), social interaction (Wacker et al., 2011; Oettl et al., 2016), controlling food intake (Soria-Gomez et al., 2014), and integrating activity within and between the two OBs (Schoenfeld and Macrides, 1984; Lei et al., 2006; Yan et al., 2008; Kikuta et al., 2010; Esquivelzeta Rabell et al., 2017; Grobman et al., 2018). Despite this wide variety of proposed functions the exact role of centrifugal AON projections in modulating ongoing OB activity remains poorly characterized. Only a few studies have investigated the influence of centrifugal AON projections on OB circuit function (Markopoulos et al., 2012; Oettl et al., 2016; Grobman et al., 2018); demonstrating that AON inputs can depolarize as well as inhibit mitral/tufted cells (MTC).

In the present study, we used optogenetic AON stimulation to decipher AON effects on odor related behavior. Whereas AON stimulation was not perceived as an odor equivalent cue, AON activation during odorant presentation reliably suppressed odor responses. This effect was constant across odors and concentrations. Optical stimulation of AON top-down projections resulted in a substantial decrease in MTC spiking frequency during sensory stimuli of different strength matching the behavioral results. These findings support the hypothesis that the AON acts as a strong regulator of olfactory information transmitted to higher brain areas.

## Materials and Methods

### Animals strain and care

We used a mouse line (Chrna7-Cre, kindly provided by S. Rogers and P. Tvrdik, University of Utah) in which an IRES-Cre bi-cistronic cassette was introduced into the 3’noncoding region of the cholinergic nicotinic receptor alpha7 (*Chrna7*) (Rogers and Gahring, 2012; Rogers et al., 2012a; Rogers et al., 2012b; Gahring et al., 2013). Animals of either sex were used. Animals were housed under standard conditions in ventilated racks. Mouse colonies were bred and maintained at RWTH Aachen University animal care facilities. Food and water were available *ad libitum* unless otherwise noted. All experimental protocols were approved by local authorities and are in compliance with European Union legislation and recommendations by the Federation of European Laboratory Animal Science.

### Viral vectors

Viral vectors were obtained from the viral vector core of the University of Pennsylvania or Addgene. Vectors were from stock batches available for general distribution. pAAV-EF1a-double floxed-hChR2(H134R)-EYFP-WPRE-HGHpA was a gift from Karl Deisseroth (Addgene viral prep # 20298-AAV1; http://n2t.net/addgene:20298; RRID: Addgene_20298). Injection of Cre-dependent vector (AAV1.EF1a.DIO.hChR2(H134R)-eYFP.WPRE.hGH (*AAV.FLEX.ChR2.YFP*)) was performed as described in (Wachowiak et al., 2013; Rothermel et al., 2014). Briefly, AON virus injection in adult (≥ 6 weeks) homozygous Chrna7-Cre mice was performed using stereotaxic targeting (relative to Bregma (in mm) +2.8 anteroposterior, 1.25 mediolateral, -2.7 dorsoventral, (Rothermel and Wachowiak, 2014)). Virus (0.5 μl; titer 1.9 × 10^12^) was delivered through a 26 gauge metal needle at a rate of 0.1 μl/min. Mice were individually housed for at least 30 days before evaluating for transgene expression or recording.

### Olfactometry

Odorants were presented as dilutions from saturated vapor in cleaned, humidified air using a custom olfactometer under computer control (Bozza et al., 2004; Verhagen et al., 2007; Rothermel et al., 2014). Odorants were typically presented for 4 - 10 seconds. All odorants were obtained at 95 - 99% purity from Sigma-Alrich and stored under nitrogen/argon. The following monomolecular odorants were used: isoamyl acetate, methyl valerate, ethyl butyrate, ethyl tiglate, sec butyl acetate, methyl caproate, 2-hexanone, 2-heptanone, cyclohexylamine, valeraldehyde, propyl butyrate, 2-pentanone, ethyl acetate, isopropyl butyrate, 4-methylanisole, 2-methylbutyraldehyde, methyl benzoate, vinyl butyrate, hexanal and Mix4 (2-hexanone, sec butyl acetate, ethyl butyrate, methyl valerate). The concentration of the odorants ranged from 0.1% to 4.5% saturated vapor (s.v.).

### Awake, head-fixed preparation

Behavioral testing in awake, head-fixed mice were adapted from previously described protocols (Wachowiak et al., 2013). For behavioral experiments, we expressed AAV.FLEX.ChR2.YFP in the AON of Chrna7 animals as previously described. A custom head bolt was affixed to the skull with its posterior edge at lambda using dental acrylic. An optic fiber (low OH, 200-μm core, 0.39 NA; FT200EMT, Thorlabs) was cut with a diamond knife and inserted into a 1.25-mm-diameter ceramic ferrule (CFLC230, Thorlabs) with a 230-μm bore. The optic fiber was adhered to the ferrule with epoxy (Epoxy 353ND Kit, Precision Fiber Products). The tip of the optic fiber was finely ground with polishing sandpaper and a grinding puck (D50-L, Thorlabs). After viral injection, the optic fiber was implanted into the targeted brain region under the guidance of a stereotactic device and a cannula holder (XCL, Thorlabs) (same coordinates as above except the fiber was positioned slightly dorsal to the injection site). To secure the implanted optic fiber, dental cement was applied to the skull surface. After complete solidification of the dental cement, the cannula holder was removed from the implanted optic fiber. In control mice viral injection was omitted. All steps were performed in a single surgical procedure under isoflurane anesthesia. Aseptic techniques were used throughout the procedure and local anesthetic (bupivacaine, 1%; Sigma-Aldrich) was applied to all incision areas.

### Behavioral testing and optical stimulation

Experiments were performed in a custom built behavioral setup. Odorant presentation, olfactometer control, water delivery, optical stimulation and data acquisition triggers were controlled with custom software written in LabView (National Instruments). All mice (n = 12, 8 ChR2 and 4 control animals) were initially trained on a simple lick/no-lick task structure. Behavioral training began 10-11 days after head bolt and optic fiber surgery. Mice were water deprived to ∼85% of baseline body weight and gradually habituated to run on a free-floating Styrofoam ball in daily sessions. Persistent limb movement or attenuated respiration was used as an indicator of stress, in which case the session was terminated. An odor delivery port was positioned ∼ 5 mm in front of the animal’s snout, and a lick spout made available for water delivery. During the initial phases of training, mice were allowed to lick for a small water reward (∼5-6 µl) at increasing intertrial intervals (ITI, 5- to 10-s, Phase I and II). After acclimation, odorants were added to the training sessions (4 s duration, Phase III). During his phase odorants (typically one per session) were passively presented and not rewarded.

Finally mice were trained in a go/no-go odor paradigm (Phase IV). Mice discriminated rewarded odorants (S+) from clean air (S-, “blank”) by licking the lick spout in response to the S+ and refraining from licking to the S-. In each behavioral session, 4 to 8 odorants (randomly picked out of a repertoire of 36 total odorants) were applied in random order. S+/S- presentation was also randomized (50/50 distribution). Odorants were presented for 4s at a concentration of 0.5% (unless stated otherwise) with an ITI varying randomly from 15 to 24 s. Incorrect licking (false alarms) at any time during presentation of the S- was punished with a 7-s increase in the following ITI. Mice were tested in a single daily session (range: 50–210 trials (each S+ or S- presentation is defined as one trial); 30–90 min). Analysis of behavioral data was performed using custom scripts in Matlab. A response on a S+ trial (hit) and no response during an S-trial (correct rejection) were categorized as correct responses during data analysis, no response on a S+ trial (miss) or a response during an S-trial (false alarm) were categorized as incorrect responses. Performance accuracy was calculated as: odor hits + correct rejections / number of trials. Animals had to reach performance accuracy > 80% before being tested further.

Mice were tested in the following paradigms: *Optogenetic stimulation in blank trials*. The go/no-go odor paradigm (Phase IV) was modified such that in ∼ 10% of S+ trials a blank was presented and licking to these trials would have been rewarded. AON photostimulation (4 sec duration) was coupled to a subpopulation of S+ blank trials. Optical stimulation trials were randomly interspersed with trials with no stimulation (S+ and S-). On average 11.5 ± 0.57 (mean ± SEM) optical stimulation trials were applied in one behavioral session. Photostimulation intensity was gradually increased from trial to trial (range 1 – 10 mW). Photostimulation light (delivered unilaterally via a 200 μm optical fiber positioned at AON; as described above) was generated by a 473 nm DPSS laser (VM-TIM). The optical stimulation protocol was adopted from a pulse protocol previously established for optical piriform cortex stimulation (Choi et al., 2011) (25 ms pulses repeated at 20 Hz for 4 sec, maximal power 10 mW). Three additional pulse protocols were tested in blank photostimulation trials: 5 ms pulses 50 Hz, 15 ms pulses 50 Hz and 10 ms pulses 20 Hz using a power between 1-13 mW. Responses in blank / blank + photostimulation trials were measured in % of total trials in this condition. Performance accuracy in this and all following paradigms was calculated as described above from non-stimulated trials and is always provided as a measure of performance and motivation of a particular animal in a task.

### Optogenetic stimulation in odor trials

The go/no-go odor paradigm (Phase IV) was modified such that AON photostimulation was coupled to a subpopulation of S+ odor trials (same parameters as above). For that, one odor was chosen and AON photostimulation was co-applied with that particular odor (4 sec duration, starting simultaneously). Selected odors changed between sessions. Optical stimulation trials were randomly interspersed with trials with no stimulation (S+ and S-). Photostimulation intensity was gradually increased from trial to trial (range 1 – 10 mW). Unless noted otherwise, all photostimulation trials before odor suppression are categorized as subthreshold stimulations. Responses in odor + subthreshold / suprathreshold photostimulation trials were measured in % of total trials in this condition.

### Optogenetic stimulation of different odors in one session

The *“Optogenetic stimulation in odor trials”* paradigm was modified so that after suprathreshold intensity was determined for one randomly chosen odorant, AON photostimulation was co-applied with different odorants within one session. Responses in odor + subthreshold / suprathreshold photostimulation trials were measured in % of total individual odorant trials.

### Optogenetic stimulation in odor concentration trials

The *“Optogenetic stimulation in odor trials”* paradigm was modified: the minimal light intensity for inhibiting odor detection at a concentration of 0.5% was determined. Using this intensity the odor concentration was gradually increased (range 0.5 – 4.5 %). The lick delay (sec, mean ± SD) of photostimulated ChR2 and control mice was determined for each tested odorant concentration (4 sec = no lick).

### Decreasing the optical stimulation length relative to odor presentation

The *“Optogenetic stimulation in odor trials”* paradigm was modified: after suprathreshold intensity was determined for one randomly chosen odorant, the overlap between laser and odor stimulation was varied (while still starting simultaneously) (2 s laser 4 s odor; 3 s laser 6 s odor; 4 s laser 6 s odor). Lick delays of ChR2 mice were compared to control animals (lick delay (sec, mean ± SD)).

### Novel odor trials

The go/no-go odor paradigm (Phase IV) was modified such that a novel odorant was applied for the first time. The number of trials until the animal started licking to this novel odorant was determined.

### Extracellular recordings and optical stimulation

MTC unit recordings and optical OB stimulation were performed as described previously with several modifications (Rothermel et al., 2014). Briefly, mice were anesthetized with pentobarbital (50 mg/kg) and placed in a stereotaxic device. Mice were double tracheotomized and an artificial inhalation paradigm used to control air and odorant inhalation independent of respiration (Wachowiak and Cohen, 2001; Spors et al., 2006; Eiting and Wachowiak, 2018). Extracellular recordings were obtained from OB units using sixteen channel electrodes (NeuroNexus, A1×16-5mm50-413-A16, Atlas Neuro, E16+R-100-S1-L6 NT) and an RZ5 digital acquisition system (TDT, Tucker Davis Technologies). Recordings from presumptive MTCs were obtained and selected as described in (Carey and Wachowiak, 2011), i.e. selected units had to be well isolated, located in the vicinity of the mitral cell layer, and show clear spiking activity in the absence of odorant. Electrode depth was monitored with a digital micromanipulator (Sutter Instruments, MP-225, or Thorlabs PT1/M-Z8). Recording sides were confined to the dorsal OB. Odorant alone (‘baseline’) and odorant plus optical stimulation trials (at least 3 trials each) were interleaved for all odorants (inter-stimulus interval 70 s). Recordings with at least 5 repeated trials of each condition were subject to unit-by-unit statistical analysis as described below.

For optical OB stimulation, light was presented as a single 10 - sec pulse either alone or simultaneous with odorant presentation using a 470 nm LED and controller (LEDD1B, Thorlabs) and a 1 mm optical fiber positioned within 3 mm of the dorsal OB surface. The light power at the tip of the fiber was between 1 and 10 mW.

### Electrophysiological data analysis

Basic processing and analysis of extracellular data followed protocols previously described for multichannel MTC recordings (Rothermel et al., 2014). Briefly, action potential waveforms with a signal-to-noise ratio of at least 4 s.d. above baseline noise were thresholded and saved to disk and single units were further isolated using offline spike sorting (OpenSorter, TDT). Responses to optical or odorant stimulation were analyzed differently depending on the experimental paradigm: stimulation effects on spontaneous spike rate (no artificial inhalation, “no-sniff” condition) were measured by calculating spikes / second (Hz) for the 9 sec before or during stimulation. Selection of ‘sniff modulated’ units was performed as described previously (Rothermel et al., 2014). Inhalation-evoked responses during inhalation of clean air (“sniff” condition) were measured by averaging the number of spikes per 1-sec period following each inhalation in the 9 inhalations pre-stimulation or during stimulation and across multiple trials (minimum of 3 trials in each condition for all units). Odorant-evoked responses were measured as changes in the mean number of spikes evoked per 1-sec inhalation cycle (Δ spikes / sniff) during odorant presentation, relative to the same number of inhalations just prior to odorant presentation. For statistical analysis, significance for changes in firing rate (baseline versus optical stimulation) was tested on a unit-by-unit basis using the Mann-Whitney *U* test on units tested with 5 or more trials per condition.

### Statistical analysis

Significance was determined using paired Student’s *t*-test, Wilcoxon signed rank test, Mann–Whitney *U* test and Kruskal-Wallis test, where appropriate. Multiple comparison test were used for post hoc comparisons. Significance was defined as *P<0.05, **P<0.005, ***P<0.0005, ****P<0.0001. All tests are clearly stated in the main text.

### Histology

Viral (AAV.FLEX.ChR2.YFP) expression in AON cells / axonal projections and optic fiber placement was evaluated with post hoc histology in all experiments to confirm accurate targeting of AON neurons and a lack of expression in OB neurons as described in (Wachowiak et al., 2013; Rothermel et al., 2014). Briefly, mice were deeply anesthetized with an overdose of sodium pentobarbital and perfused with 4% paraformaldehyde in PBS. Tissue sections were evaluated from native fluorescence without immunohistochemical amplification with a Leica TCS SP2 confocal laser scanning microscope at 10x or 20x magnification.

## Results

We used a mouse line in which an IRES-Cre bi-cistronic cassette was introduced into the 3’ noncoding region of the cholinergic nicotinic receptor alpha 7 gene (Chrna7) (Rogers et al., 2012b), as its expression is high in AON neurons (Dominguez del Toro et al., 1994; Brunjes et al., 2005), to target cortical top-down projections from the AON to the OB (Rothermel and Wachowiak, 2014). We expressed ChR2(H134R)-EYFP selectively in AON neurons using a Cre-dependent viral expression vector targeted to the AON by stereotaxic injection. Viral injection centrally into the AON resulted in a robust cellular ChR2-EYFP expression in all major AON subdivisions (pars principalis, dorsal, lateral, medial and ventral part; including pars externa) (Figure 1A). Labeled cells displayed one or more thick apical dendrites typical of pyramidal neurons in agreement with previous description of AON projections neurons (Brunjes and Kenerson, 2010; Rothermel and Wachowiak, 2014). Four weeks following unilateral virus infection, ChR2-EYFP protein was apparent in AON fibers throughout the OB (Figure 1B): ipsilateral AON axon terminals targeted mainly the granule cell layer (GCL) and, to a lesser extent, the external plexiform layer (EPL), whereas fewer fibers in the contralateral EPL could be observed (Figure 1B), consistent with earlier reports (Reyher et al., 1988; Rothermel and Wachowiak, 2014). Based on these results we conclude that our viral approach mainly labels AON pars principalis neurons reported to have bilateral OB projections (Brunjes et al., 2005; Wen et al., 2019).

**Figure 1.**
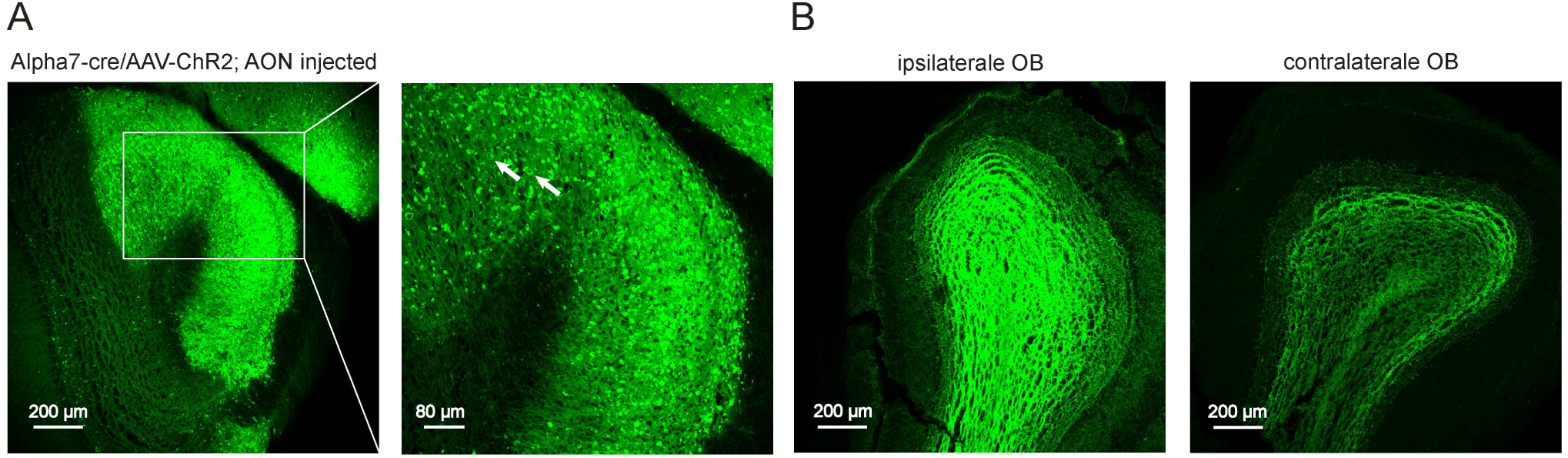
ChR2 expression after unilateral viral injection into AON of a Chrna7-Cre mouse. A. Left: Low-magnification confocal stack showing heavy ChR2-EYFP expression in AON neurons in all AON subdivisions (2.73 bregma; coronal section). Right: Magnification of the lateral AON region showing strong cellular ChR2-EYFP expression. White arrows indicate exemplary positive labelled AON neurons. B. Left: ChR2-EYFP-expressing AON fibers in the ipsilateral OB imaged 4 weeks after AAV-ChR2-EYFP injection into the AON of a Chrna7-Cre mouse. ChR2-EYFP labelled AON axon terminals targeted primarily the GCL and the EPL. Right: Confocal stack from the contralateral OB displaying AON terminals predominantly in the GCL. EPL: external plexiform layer; MCL: mitral cell layer; GCL: granule cell layer.

### Activation of AON neurons in awake animals is not perceived as an odorant equivalent cue

In order to explore possible behavioral effects of AON activation in odorant perception, we initially asked whether photostimulation of ChR2-expressing neurons in the AON could serve as an odorant equivalent cue. The AON as a sensory olfactory cortical area receives direct mitral / tufted cell input from the OB. Since optogenetically induced “illusory” sensory perception has been reported for different sensory areas (O’Connor et al., 2013; Guo et al., 2015) we tested if AON activation might be able to evoke odor sensations in mice. We used a go/no-go odor discrimination assay in which water-restricted head-fixed mice freely running on a Styrofoam ball (Figure 2A) were exposed to different odors (S+), all of which were rewarded (Figure 2B). In each behavioral session, 4 to 8 odorants (randomly picked out of a repertoire of 36 total odorants) were applied in random order (50–210 trials (S+ and S-presentations) per session; 30–90 min duration). S+/S- presentation was also randomized (50/50 distribution). During this period, mice learned to lick to any given odor stimulus and to refrain from licking to clean air (S-, blank trial). When mice performed reliably above criterion (> 80% accuracy; odor hits + correct rejections / number of trials, see Methods) we tested whether unilateral photostimulation of ChR2 expressing AON neurons would elicit licking responses when applied without sensory input (3 ChR2 and 3 controls, 27 sessions). Therefore, we modified the behavioral paradigm such that in ∼ 10% of S+ trials we presented a blank (clean air) instead of an odorant (gray bars in the S+ condition, Figure 2B) and licking to these trials would have been rewarded. As expected, mice did not lick to S+ blank trials (Figure 2B, C). When we coupled the AON photostimulation to a subpopulation of S+ blank trials (grey bars with blue surrounding) mice equally failed to respond suggesting that activating AON was not able to elicit an odor percept. In order to ask if cortical AON activation might trigger odor perception, this low number of optogenetic stimulation trials was necessary as mice had to stay trained to associate water reward with real odor presentation. Importantly, mice were not trained to report to optogenetic stimulation since it has been shown that with sufficient training (≥ 650 stimulations) animals can be trained to report essentially any kind of rewarded stimulus in any brain area e.g. from optogenetic stimulation of one olfactory bulb glomerulus (Smear et al., 2013) to electrical activation of single cells (Histed et al., 2013). Responses in blank trials did not differ significantly from responses in blank + photostimulation trials within and between individual ChR2 or control animals (Figure 2C, % blank responses 2.22 ± 1.28 control, 4.1 ± 2.4 ChR2; % blank + photostimulation responses 0.83 ± 0.48 control, 0 ChR2; *p* = 0.42, Kruskal-Wallis test). We performed the experiments using four different optogenetic pulse protocols. The absence of a licking response in the photostimulation trials was not a result of missing motivation as general performance accuracy (odor hits + correct rejections / number of trials) was high and not significantly different within and between individual ChR2 and control mice (93.24 ± 1.9 % control, 92.9 ± 0.34 % ChR2 (mean ± sem); *p* = 0.14, Kruskal-Wallis test). Mice also failed to respond to AON stimulation when this paradigm was repeated for six consecutive days, showing that mice did not report optogenetic stimulation with this amount of stimulation trials.

**Figure 2.**
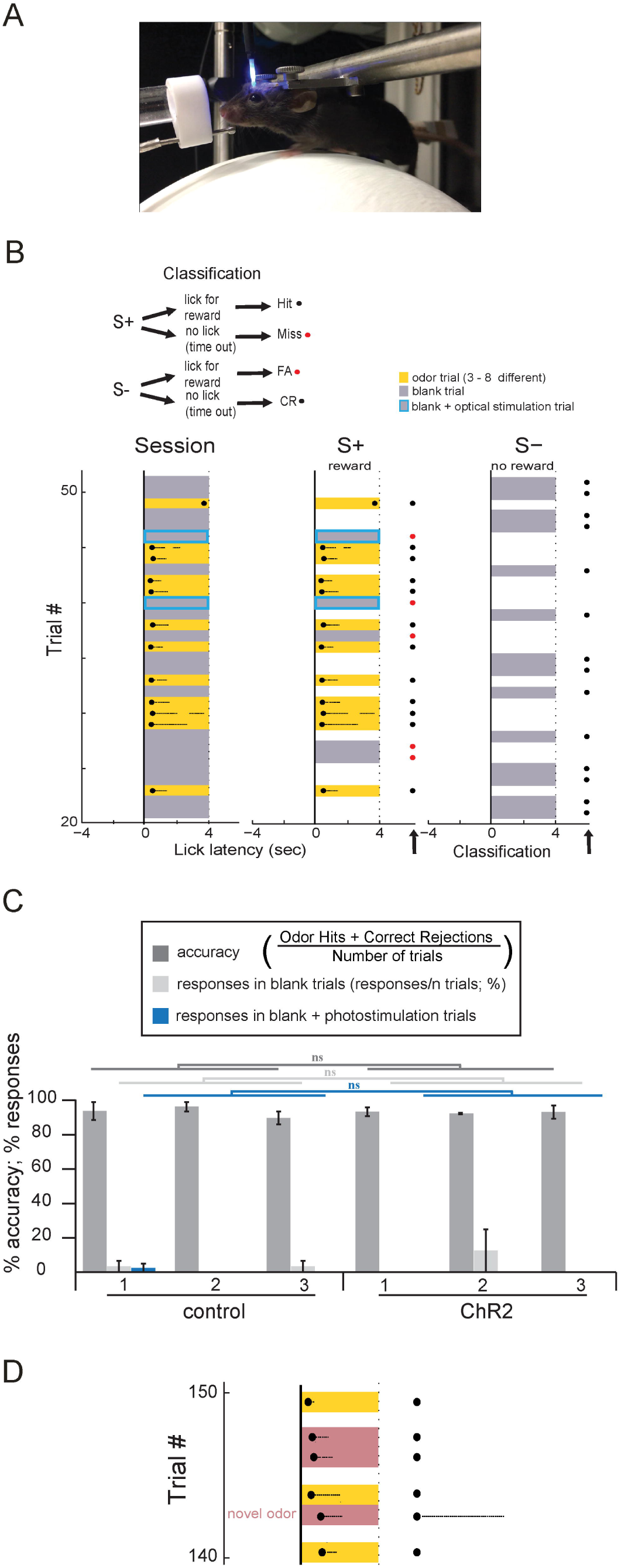
Optogenetic AON activation is not perceived as an odorant equivalent cue. A. Behavioral setup depicting head-fixed mouse on a freely floating Styrofoam ball. B. Representative part of one behavioral session (left, ordinate displays trial number in chronological order). Mice were trained in a go/no-go odor paradigm and discriminated rewarded odorants (S+, yellow squares, odor duration 4 seconds) from clean air (S-, “blank”, gray squares) by licking in response to the S+ and refraining from licking to the S-. 3 to 8 odorants (randomly picked out of a repertoire of 36 total odorants) were applied in random order. For simplification all odorants in this session are depicted as yellow squares. Middle and right panels represent behavioral session sub-trials ordered by S+ (middle) and S-(left) trials. First licks in each trial are represented by solid black dot; subsequent licks as lighter dots. Each trial is categorized according to the classification depicted on top. Briefly, response on a S+ trial (hit) and no response during an S-trial (correct rejection) were categorized as correct responses (black dots at the end of the odor presentation); no response on a S+ trial (miss) or a response during an S-trial (false alarm) were categorized as incorrect responses (red dots at the end of the odor presentation). ∼ 10% of S+ trials consisted of a blank presentation and licking to these trials would have been rewarded. Blank + AON photostimulation trials (in S+ condition) are highlighted by a blue box. Mice did not lick to S+ blank or S+ blank+photostimulation trials. C. Odor response accuracy and % responses in blank and blank+photostimulation trials for control and ChR2 animals. No significant difference was observed within and between individual ChR2 or control mice (n = 6 mice, 27 sessions). D. Part of a representative “*Novel odor trial”* session. The animal responded to the novel odorant in the first trial.

Next, we asked if the lack of responses in blank + photostimulation trials might be due to a failure of the mice to generalize from the 36 different odors they had been trained to. In this case mice might perceive AON stimulation as an odorant but fail to respond due to its novelty. In order to test if animals generalized to different, novel odorants we introduced novel odorants that were never encountered before in a subset of sessions (4 ChR2 animals, 10 sessions, Figure 2D). In 8 sessions ChR2 mice licked directly to the novel odor, while in 2 sessions it took only two trials to lick to the presentation of this novel odor. Control animals also directly responded to the presentation of a novel odorant (3 controls, 6 sessions). This instantaneous licking response to a novel odor stimulus argues that animals generalized to odors and therefore renders it unlikely that an AON stimulation by itself (in the blank condition) caused a different/ novel odor percept that was not recognized. In conclusion these data suggest that cortical AON activation does not seem to trigger odor perception.

### AON activation impairs odor detection in awake behaving animals

Next, we asked whether a photostimulation of ChR2-expressing neurons in the AON might affect odor processing. Experiments were partially conducted in the same animal cohort used for the previous behavioral experiments. A similar go/no-go odor discrimination assay was used in which mice were trained to lick to different odors (S+), and refrain from licking to clean air (S-). Each behavioral session consisted of an initial baseline session (duration range from 10 –30 minutes; 20 -100 trials) in which the mouse’s response to different odors (S+) and clean air (S-) was determined. Both experimental groups performed reliably above criterion (80% accuracy). Following this baseline period, one odor was randomly chosen and AON photostimulation was co-applied with that odor (Figure 3A; (4 sec duration, starting simultaneously)) (20 Hz with 25 ms pulses, for 4 s; adopted from (Choi et al., 2011)). Odor + AON stimulation trials were interleaved with S- and odor only trials (Figure 3A; 8 ChR2 and 4 controls, 130 sessions). During this period the photostimulation intensity was gradually increased (range 1 – 10 mW). Plotting lick latency as a function of laser power (Figure 3B) revealed a photostimulation power-dependent increase in lick latency (0.51 ± 0.03 to 1.5 ± 0.31 sec; photostimulated odor, *p* = 0.03, Wilcoxon signed rank test) that was restricted to the odor coupled to photostimulation in the ChR2 group (0.46 ± 0.02 to 0.56 ± 0.11 sec; non-photostimulated odors, *p* = 0.36, Wilcoxon signed rank test). A further slight increase in photostimulation intensity finally caused a failure of odor responses (0.51 ± 0.03 to 4 sec (licktimeout); photostimulated odor, *p* = 0.03, Wilcoxon signed rank test) of only that particular odor coupled to light stimulation (0.46 ± 0.02 to 0.58 ± 0.1 sec; non-photostimulated odors, *p* = 0.46, Wilcoxon signed rank test, Figure 3B). This effect was not observed in control animals (0.45 ± 0.01 to 0.43 ± 0.02 sec; photostimulated odor, *p* = 0.44; 0.45 ± 0.02 to 0.45 ± 0.01 sec; non-photostimulated odors, *p* = 0.9, Wilcoxon signed rank test). “Suprathreshold” photostimulation significantly inhibited responses to odor presentation (measured in % of the total number of trials in this condition) in all ChR2 mice when compared to control animals (Figure 3C, blue bars, 96.99 ± 1,9 % control, 1,93 ± 0,66 % ChR2 (mean ± sem); *p* = 5 × 10^−10^, Kruskal-Wallis test, NP posthoc test). The reduction of a licking response in these “suprathreshold” photostimulation trials was however not a result of a declining motivation as general performance accuracy as well as percentage of responses during odor presentation in subthreshold stimulation trials were not statistically different within and between ChR2 and control mice (Figure 3C, dark and light gray bars, accuracy: 96.29 ± 0.27 % control, 91.1 ± 0.76 % ChR2; *p* = 0.19, subthreshold stimulation: 96.45 ± 1.21 % control, 92.74 ± 1.67 % ChR2; *p* = 0.13 Kruskal-Wallis test). Additionally, odor presentation without photostimulation led to a normal response in ChR2 mice. Control mice did not show any effects upon photostimulation regardless of the applied laser power (range 1 – 10 mW) as did ChR2 mice when trained to a non-olfactory detection task (data not shown), showing that mice could perform well during light stimulation

**Figure 3.**
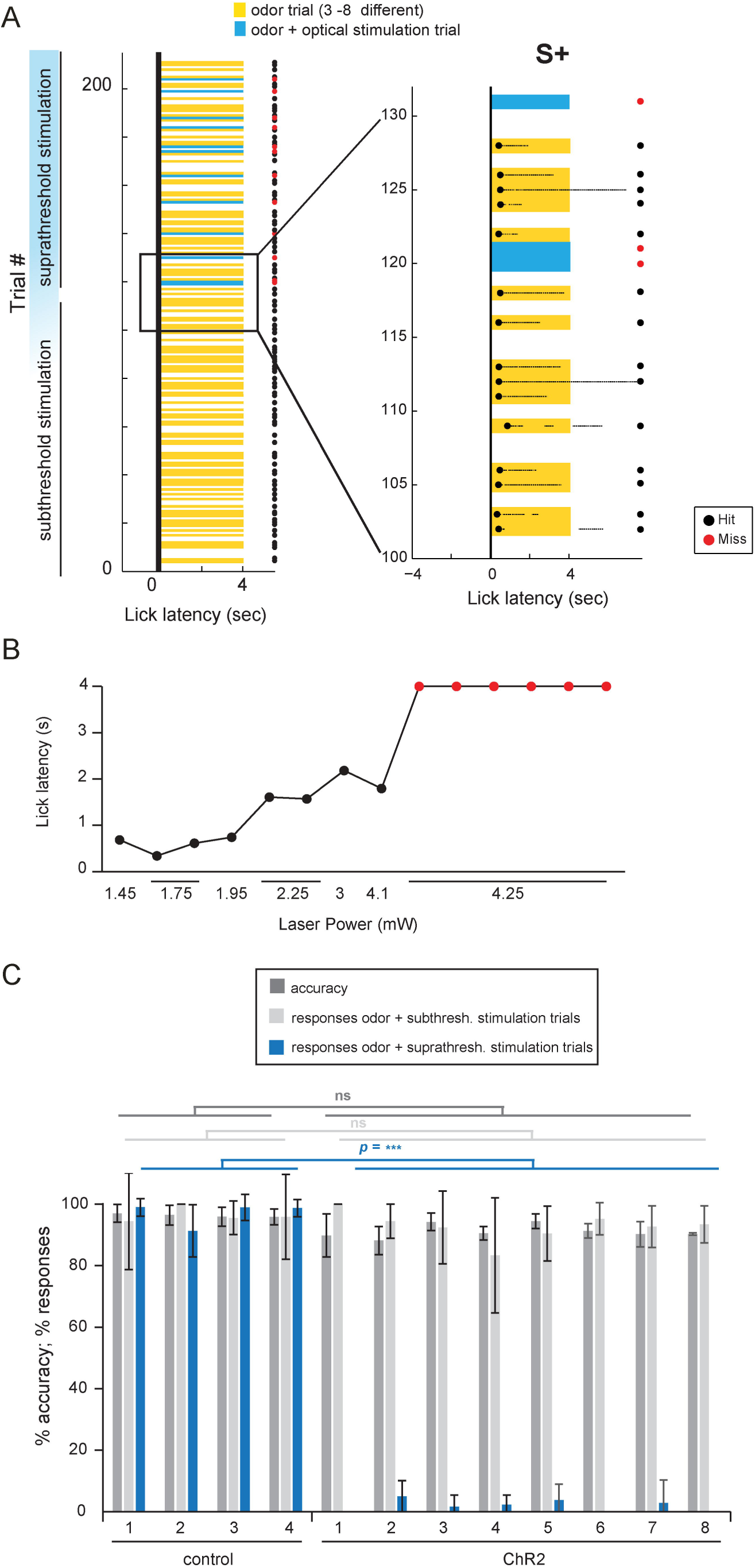
Optogenetic AON activation suppresses odorant detection. A. Representative go/no-go behavioral session. Mice were trained in a go/no-go odor paradigm and discriminated rewarded odorants (S+, yellow squares, odor duration 4 seconds) from clean air (S-, “blank”, gray squares) by licking in response to the S+ and refraining from licking to the S-. 3 to 8 odorants (randomly picked out of a repertoire of 36 total odorants) were applied in random order. For simplification all odorants in this session are depicted as yellow squares. One odor was chosen and AON photostimulation was co-applied with that particular odor (4 sec duration, starting simultaneously). Selected odors changed between sessions. Optical stimulation trials were randomly interspersed with trials with no stimulation (S+ and S-). Photostimulation intensity was gradually increased from trial to trial (range 1 – 10 mW). Unless noted otherwise, all photostimulation trials before odor suppression are categorized as subthreshold stimulations (suprathreshold trials are depicted in blue). Lick responses are only depicted in the magnification on the right. AON photostimulation strongly suppresses odorant detection. B. Lick latency (s) plotted as a function of laser power (mW). Increased laser power led first to an increase in lick latency before causing complete suppression of odorant evoked licking. C. Odor response accuracy and % responses during subthreshold / suprathreshold odor + photostimulation trials for control and ChR2 animals. Suprathreshold stimulation selectively suppresses odorant detection in all tested ChR2 mice. There was no difference in task accuracy and % responses in subthreshold trials within and between control and ChR2 mice (n = 12 mice, 130 sessions).

We also tested if AON photostimulation is able to suppress responses to different odorants. Therefore, in separated sessions different odorants were coupled to the photostimulation. Licking responses could not only be inhibited by AON photostimulation for all tested monomolecular odorants but also for an odor mixture (Table 4-1, 8 ChR2 animals, 97 sessions, 12 monomolecular odorants, one mixture of four odorants; each odorant was tested in a least 4 session). Lick responses to different odorants could also be inhibited within a single behavior session by quickly switching optical stimulation (Figure 4A) between up to 6 odorants. This demonstrates that 1) AON mediated inhibition of odor responses is not odor specific 2) the AON inhibits odor responses on a fast timescale and 3) there is no training effect involved. As shown previously, animals were able to generalize to a novel odor stimulus (Figure 2D) which renders it unlikely that the absence of licking in odor + photostimulation trials is caused by an AON mediated change in/or novel odor percept. Photostimulation in control animals had no significant effect on odor responses (Table 4-1 A, 4 controls, 101 sessions, 12 monomolecular odorants, one mixture of four odorants). We also examined whether the effect of AON activation could be overcome by stronger sensory input. For this, we set the laser strength for optical stimulation at the minimal light intensity necessary for the inhibition of odor detection at a concentration of 0.5%. Odor concentration was then gradually increased from 0.5% to 4.5% (Figure 4B). AON photostimulation significantly inhibited odor detection over this whole concentration range (lick delay (sec, mean ± SD) of ChR2 mice (4 animals, 40 sessions) vs. photostimulated controls (3 animals, 53 sessions); 4 sec = no lick, 0.1%, 4 ± 0 sec vs. 0.52 ± 0,05 sec, *p* =5.8 × 10^−4^; 0.5%, 3.97 ± 0.06 sec vs. 0.58 ± 0.57 sec, *p* =2.5 × 10^−4^; 1%, 4 ± 0 sec vs. 0.83 ± 0.91 sec, *p* =1 × 10^−4^; 2%, 4 ± 0 sec vs. 0.4 ± 0.08 sec, *p* = 4.6 × 10^−5^; 3%, 4 ± 0 sec vs. 0.4 ± 0.05 sec, *p* = 4.5 × 10^−5^; 4%, 3.81 ± 0.36 sec vs. 0.56 ± 0.32 sec, *p* = 0.009; Mann–Whitney *U* test for ≥ 4 session per odorant concentration).

**Table 4-1:** Optogenetic AON activation inhibits odorant detection independent of odor identity. AON photostimulation was co-applied with different odorants within one session. Photostimulation significantly reduces licking responses in suprathreshold trials across all tested odorants (12 monomolecular odorants, one mixture of four odorants) in ChR2 mice (8 animals, 97 session). No significant difference in performance between subthreshold and suprathreshold trials was observed in control animals (4 animals, 101 sessions).

| Odor | # of sessions | control |  | *p* value | ChR2 |  | *p* value |
| --- | --- | --- | --- | --- | --- | --- | --- |
|  |  | subthresh. | suprathresh. |  | subthresh. | suprathresh. |  |
| IAA | 23 | 93.52 | 98.56 | ns | 93.43 | 2.00 | *** |
| MV | 29 | 100.00 | 94.82 | ns | 94.72 | 0.00 | *** |
| ET | 10 | 91.67 | 100.00 | ns | 94.44 | 0.00 | * |
| PB | 14 | 99.60 | 97.33 | ns | 97.22 | 5.00 | * |
| MH | 15 | 100.00 | 97.27 | ns | 89.32 | 1.79 | *** |
| secBA | 19 | 100.00 | 94.65 | ns | 91.51 | 0.67 | *** |
| EB | 16 | 100.00 | 96.75 | ns | 92.61 | 0.00 | *** |
| MB | 12 | 100.00 | 100.00 | ns | 90.29 | 2.22 | ** |
| 2-Hep | 13 | 99.24 | 94.84 | ns | 94.45 | 0.00 | *** |
| 2-Hex | 10 | 90.28 | 98.72 | ns | 95.00 | 0.00 | * |
| VB | 9 | 95.05 | 100.00 | ns | 94.33 | 6.25 | * |
| 2-Pen | 17 | 100.00 | 100.00 | ns | 92.51 | 2.38 | *** |
| Mix4 | 11 | 95.42 | 98.57 | ns | 98.75 | 0.00 | * |

**Figure 4.**
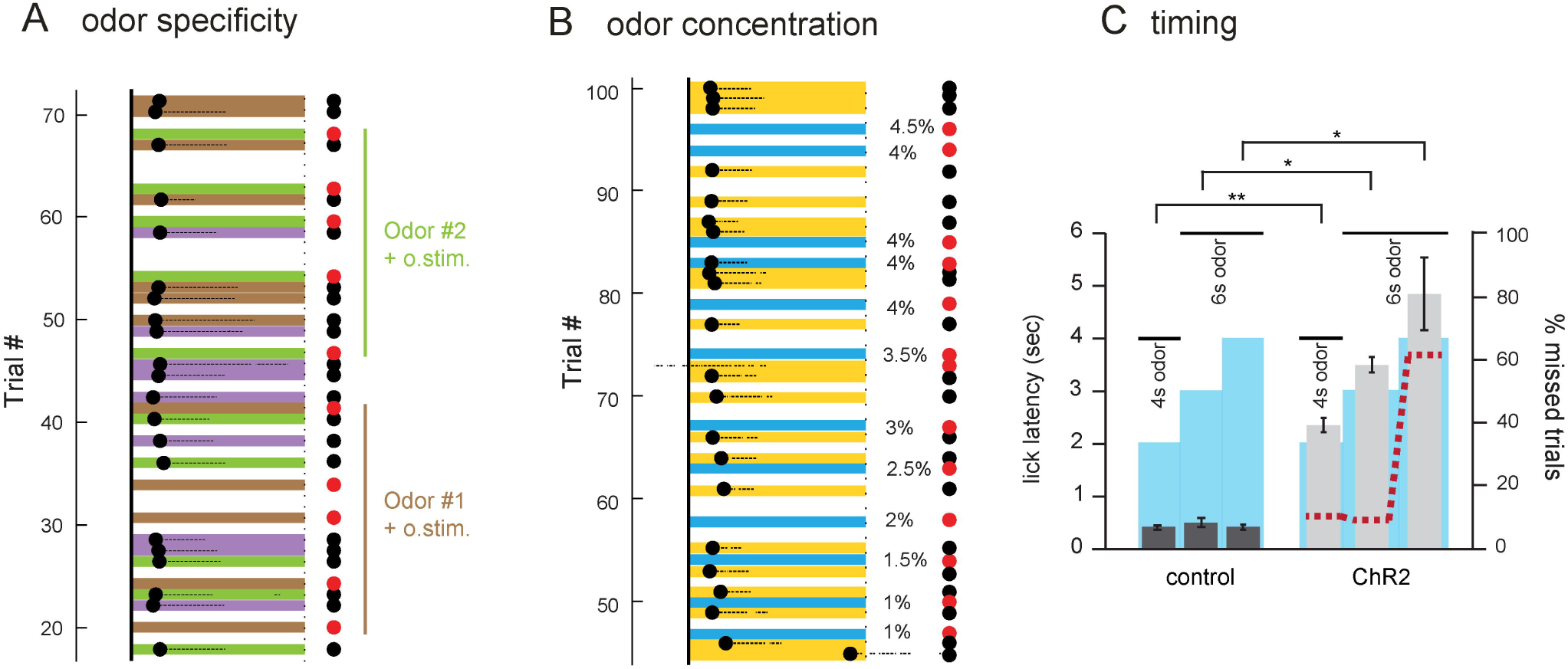
Optogenetic AON activation inhibits odorant detection independent of odor identity and concentration. A. Lick responses to different odorants could also be inhibited within a single behavior session by quickly switching optical stimulation between odorants. Representative part of one behavioral session showing suppression of odor detection for two different odorants. In contrast to previous plots, individual odorants are color coded. B. AON photostimulation significantly inhibits odor detection over a large concentration range. Representative part of a training session in which odor concentration was gradually increased from 1-4.5 %. Suppression of odor detection occurred at all tested concentrations. C. Timing of AON photostimulation affects behavioral responses. Decreasing the photostimulation time relative to the odor duration lead to an immediate response (lick latency (s)) at the end of the photostimulation in ChR2 animals. Photostimulation duration is indicated by the blue box; odor stimulation length by the black bar. Longer stimulation times also increased the number of missed trials (red dotted line).

Finally, we investigated the relative timing of AON photostimulation effects on behavioral responses by varying stimulation times from the default settings (4 sec both, given simultaneously). Using a paradigm of decreased photostimulation time relative to the odor duration, we found that ChR2 mice (3 animals, 12 sessions) immediately licked at the end of the photostimulation: a 2, 3 and 4 second optical stimulation that was followed by several seconds of “unmasked” odor, resulted in a lick delay of 2.38 ± 0.08, 3.52 ± 0.08 and 4.87 ± 0.22 seconds, respectively (Figure 4C). Lick delays of ChR2 mice at any photostimulation length were significantly different compared to control mice (3 animals, 17 sessions) (lick delay (sec, mean ± SD) of ChR2 mice vs. photostimulated controls; 2 sec photostimulation 2.38 ± 0.08 vs 0.42 ± 0.07 sec *p* = 0.004; 3 sec photostimulation 3.52 ± 0.08 vs 0.46 ± 0.06 sec *p* = 0.02; 4 sec photostimulation 4.87 ± 0.22 vs 0.43 ± 0.04 sec *p* = 0.03; Mann–Whitney *U* test). Interestingly, ChR2 mice failed significantly more often to lick in response to 4 second photostimulation trials compared to shorter stimulation lengths (Figure 4C, % missed trials; 2 sec photostimulation 9%, 3 sec photostimulation 7%, 4 sec photostimulation 61%; *p* = 2.5 × 10^−9^, Kruskal-Wallis test, NP posthoc test). This suggests that longer photostimulation produces longer lasting effects. In contrast to that, varying the odor length at a fixed photostimulation, had no significant effect on lick latency (laser 2 sec, lick delay: 2.7 ± 0.18 sec (4 sec odor), 3.07 ± 0.24 sec (6 sec odor), 2.5 ± 0.11 sec (8 sec odor), *p* = 0.12, Kruskal-Wallis test; data not shown).

Taken together we found that in mice trained to associate water rewards with odorants, optogenetic AON stimulation alone was not able to elicit a behavioral response. AON stimulation during odor presentation however, strongly suppressed licking responses. Since the AON sends strong top-down projections to the first relay station of olfactory information processing we next tested for direct AON mediated modulation of OB output activity.

### Optogenetic activation of AON axons in the OB has an inhibitory effect on spontaneous and inhalation-evoked MTC spiking

In order to investigate potential cellular correlates for the observed behavioral effects we looked at the first relay station of olfactory information in the brain, the olfactory bulb which receives extensive projections from the AON. For this we directed 473 nm light (at similar power levels used in behavioral experiments; 1-10 mW total power) onto the dorsal OB surface while recording multi-channel electrical activity from dorsally located presumptive MTCs in anesthetized, double-tracheotomized mice (Figure 5H) (see Materials and Methods).

**Figure 5.**
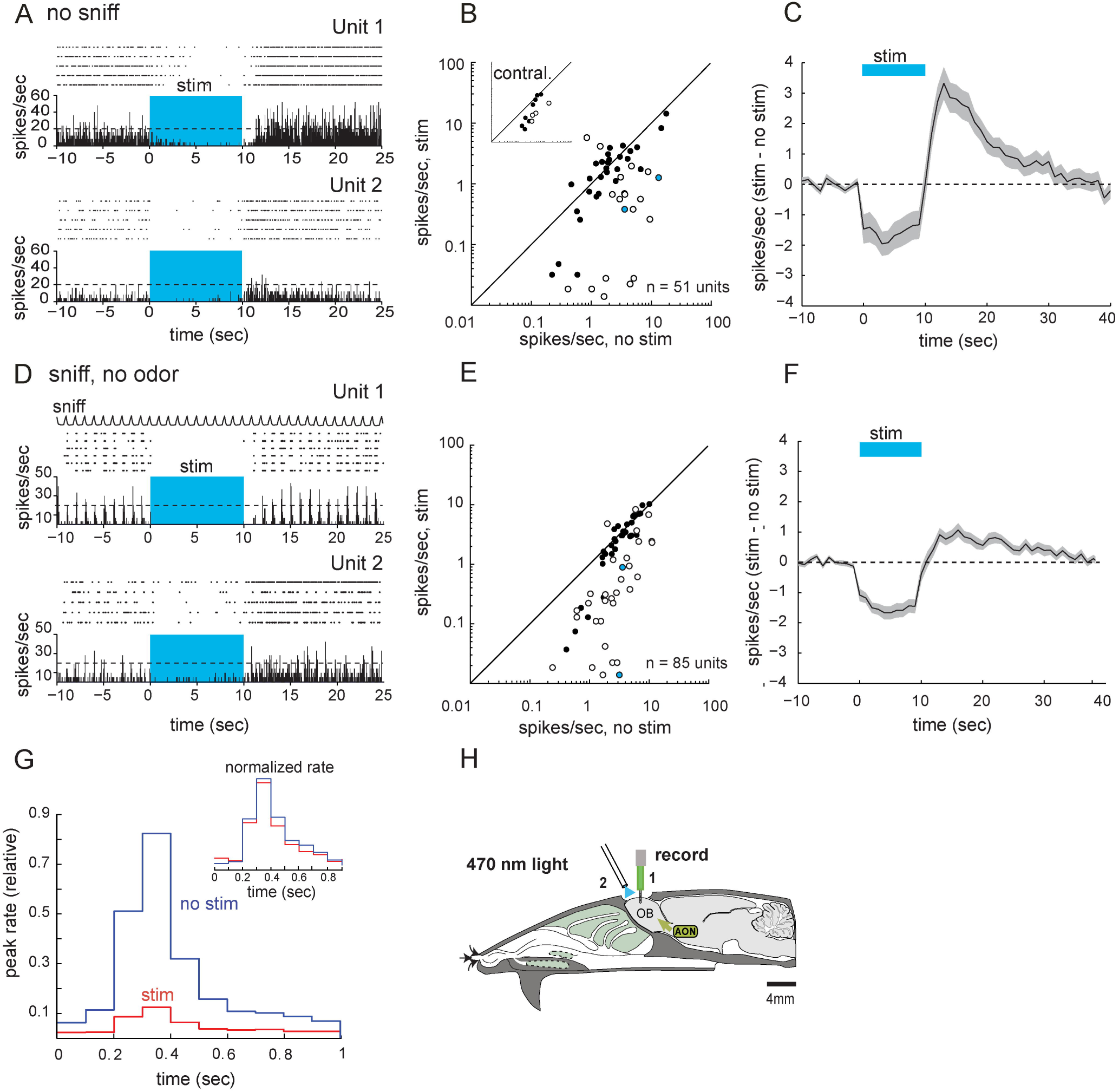
Optogenetic activation of AON axons in the OB inhibits spontaneous and inhalation-evoked MTC spiking. A. Raster plot and spike histogram of spontaneous spiking of two presumptive MTCs in the absence of inhalation (no sniff condition). The spike rate was calculated per 100 ms bin. Spike rate decrease during optical stimulation of the dorsal OB (“stim”, blue shaded box). B. Plot of spontaneous MTC firing activity in the 9 s prior (“no stim”) and during light stimulation (“stim”) for all tested units (n= 51, 8 mice). Firing activity of most units was reduced during photostimulation. All units were subjected to a unit-by-unit test for significant effects of AON stimulation and open circles indicate units that showed significant light-evoked changes. 9 showed a statistically significant reduction in firing rate and only two showed an increase. Insert: Spiking activity was also reduced when exclusively stimulating AON derived fibers in the contralateral OB in unilateral injected animals (n = 13 units from two mice). C. Time course of light-evoked (blue bar) changes in spontaneous MTC firing (mean ± SEM across all units). The trace indicates change in mean spike rate in 1 s bins relative to the mean rate before stimulation. The time axis is relative to time of stimulation onset. A clear reduction in spiking activity can be observed during optical stimulation. D. Raster plot and spike histogram of MTC spiking during inhalation of clean air and optical stimulation (blue shaded area). Inhalation-evoked spike rates decreased during optical stimulation. E. Plot of inhalation-evoked spontaneous MTC spiking in the 9 s prior (“no stim”) and during light stimulation (“stim”) for all tested units (n= 85, 10 mice). Firing activity of most units was reduced during photostimulation. Open circles depict units that showed significant light-evoked changes in firing activity when tested on a unit-by-unit basis. 52 units showed a significant decrease in spikes per sniff, only 2 units showed an increase. F. Time course of stimulation-evoked changes in inhalation-evoked MTC spiking. Spiking activity is strongly reduced during optical stimulation. G. Sniff-triggered spike histogram of inhalation-evoked MTC spiking aligned to the start of inhalation of clean air before (blue) and during (red) optical stimulation, normalized to the maximum bin in the no-stimulation condition. Bin width, 100 ms. The histogram is compiled from significant units, with firing rate normalized separately for each unit. Optical stimulation strongly reduced peak spike rate. Inset, Sniff-triggered spike histogram normalized to the maximum and minimum bin for the two conditions independently. No change in spiking dynamics after OB stimulation were observed. H. Schematic of experimental approach. See Materials and Methods for details. (1) 16-channel silicon probe (2) optical fiber. Courtesy of A.C. Puche, modified from (Aungst et al., 2003).

First, to assess the impact of AON fiber stimulation on MTC excitability in the absence of sensory input, we optically activated AON axons without ongoing inhalation (Figure 5A, B and C). In the absence of inhalation derived input, MTCs display irregular firing (Carey and Wachowiak, 2011; Courtiol et al., 2011; Rothermel et al., 2014). Optical stimulation led to a significant reduction in MTC spontaneous spiking, from 3.4 ± 3.68 Hz (mean ± SD) before stimulation, to 1.81 ± 2.45 Hz during stimulation (n = 51 units from 8 mice; *p* = 1.52 × 10^−4^, Wilcoxon signed rank test). Only cells in which the stimulation was repeated ≥ 5 times were included for analysis. This criterion supports a test of significance on each unit; 21 of these units (41%) showed a significant stimulation-evoked change in firing activity when tested on a unit-by-unit basis (Mann–Whitney *U* test). Out of these cells, 19 showed a statistically significant reduction in firing rate and only two showed an increase (Figure 5B). Among these 19 units, the median firing rate decreased by 2.89 ± 3.06 Hz. In all recorded units the decrease in spontaneous firing rates persisted for the whole duration of the optical stimulation (10 sec). Following the stimulation, an increase in spiking was observed that returned to prestimulus levels within 25 s after stimulation ceased (Figure 5A, C). Spiking activity was also reduced when exclusively stimulating AON derived fibers in the contralateral OB in unilateral injected animals (n = 13 units from two mice; 3.16±2.31 Hz before stimulation, 2.16±2.05 Hz during stimulation; *p*=0.017, Wilcoxon signed rank test, Figure 5B insert). In control mice, the same optical stimulation protocol failed to significantly modulate MTC spontaneous spiking (n= 15 units from two mice; 7.43±3.51 Hz before stimulation, 7.51±3.65 during stimulation; *p*= 0.5707, Wilcoxon signed rank test; data not shown). Thus, optogenetic activation of AON fibers leads to a strong reduction of spontaneous spiking of MTCs.

Next, we investigated the effect of AON axon activation on MTC responses during artificial inhalation of clean air. Inhalation-linked spiking pattern could be observed in 85 units from 10 mice most likely reflecting weak sensory-evoked responses (Grosmaitre et al., 2007; Carey et al., 2009; Rothermel et al., 2014). Optical stimulation of AON axons strongly diminished inhalation-linked spiking of MTCs (Figure 5D). However, an initial but fast decaying (within 30 ms) excitatory stimulation effect (Figure 5-1) similar to that reported by (Markopoulos et al., 2012) could be observed by adjusting the bin size from 100 ms to1 ms. Light–induced reduction of spiking activity was highly significant across the population of presumptive MTCs, with median spike rate decreasing from 3.07 ± 2.4 Hz to 1.66 ±2.39 Hz during optical stimulation (p = 7.45 × 10^−11^, Wilcoxon signed rank test). When tested on a unit-by-unit basis, 54 of 85 recorded units (62.35%) showed a significant optical stimulation evoked change in spiking (Figure 5E). 52 units showed a significant decrease in spikes per sniff. Inhalation evoked spiking of these 52 cells decreased by 2.2*±* 1.69 spikes/ sniff/s. Only 2 units showed an increase in spikes per sniff. Sniff-triggered spike histograms depicting MTC spiking within the course of one inhalation/sniff showed that although AON axon stimulation in the OB strongly reduced peak spike rate, it did not alter the temporal pattern of MTC responses relative to inhalation (Figure 5G) i.e. the time bin of peak firing did not change across the population of recorded units (*p* = 0.16, paired *t* test comparing time bin of the peak of inhalation-evoked firing rate for baseline vs optical stimulation). Across the population of all recorded units, the decrease in inhalation-evoked spike rates persisted for the duration of the 10 s optical stimulation (Figure 5F) and was not significantly different from reduction in spike rate observed on spontaneous MTC firing (decrease by 1.59 ± 2.94 spontaneous MTC spiking condition, 1.41 ± 1.82 inhalation evoked spiking condition, *p* = 0.46 Mann–Whitney *U* test). Following stimulation, spike rate increased above baseline levels for <25s before returning to prestimulus levels (Figure 5D Unit 2, Figure 5F). Thus, transient activation of AON axons in the OB leads to a reduction in inhalation linked MTC activity without grossly reorganizing temporal patterns of sensory input. This inhibition has a fast onset and persists for the duration of optical stimulation.

**Figure 5-1.**
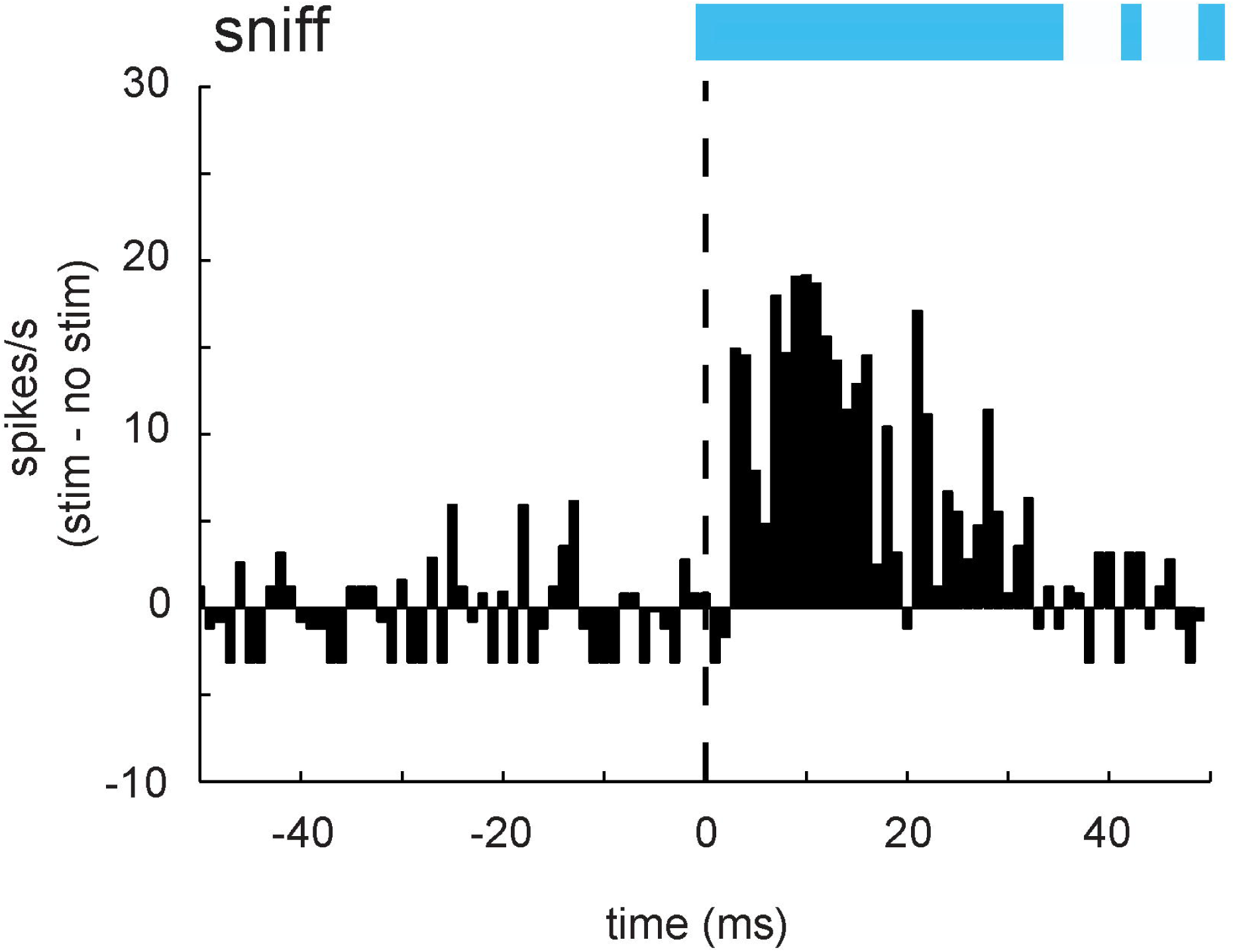
Optogenetic AON stimulation elicits fast but brief excitatory responses in MTC. Time course of changes in spontaneous firing rate 50ms before and 50ms after the start of the optical stimulation during ongoing inhalation (sniff). The mean firing rate of all recorded units (n= 85 units, 10 mice) during the 1ms time bins is depicted as spikes/s (stim - no stim). In the first milliseconds of optical stimulation MTCs show an increase in spontaneous spiking.

### Optogenetic activation of AON axons in the OB suppresses odorant-evoked MTC spiking

Since odor detection was strongly suppressed in our behavioral data we next evaluated the impact of bulbar AON modulation on odorant responses by comparing MTC odorant responses with and without optogenetic AON activation (Figure 6A). Across the population of recorded presumptive MTCs (n= 55 units, from 6 mice), AON axon stimulation significantly decreased MTC odor activity (Figure 6A, B, and C), with a decrease from 4.02 ± 2.37 spikes/sniff/s (median ± SD) during odorant presentation alone to 2.95 ± 2.96 spikes/sniff/s during odorant paired with light (*p*=1.55 × 10 ^-4^, Wilcoxon signed rank test). Optical stimulation also decreased the odorant-evoked component of the MTC response when evaluated on a single cell level (measured as Δ spikes/sniff/s relative to preodor presentation (Figure 6B). Tested on a unit-by-unit basis, 19 out of the 55 recorded cells (35%) showed a significant change in odor-evoked spiking. 17 of those cells showed a significant decrease, only two showed an increase. The median decrease in spike rate, across the 17 cells showing a significant decrease, was 2.64 spikes/sniff/s (from 3.42 ± 1.84 to 0.78 ± 1.15). As with effects on spontaneous spiking, the reduction of odorant-evoked MTC responses occurred rapidly and persisted for the duration of optical stimulation. The light mediated inhibition was followed by an increase in spiking that lasted for <25 s after the end of optical stimulation (Figure 6A, Unit 1 and 2; Figure 6C). As with inhalation-evoked MTC responses, no change in the temporal component of MTC odor responses could be observed (Figure 6D, inset, *p* = 0.6, paired *t* test comparing time bin of the peak of odorant-evoked firing rate for baseline vs optical stimulation; n = 55 units from 6 mice). Optical stimulation resulted in a uniform spike reduction across the sniff cycle (Figure 6D). Overall, this shows that activating AON axons not only strongly decreases MTC spontaneous activity but also sensory evoked responses.

**Figure 6.**
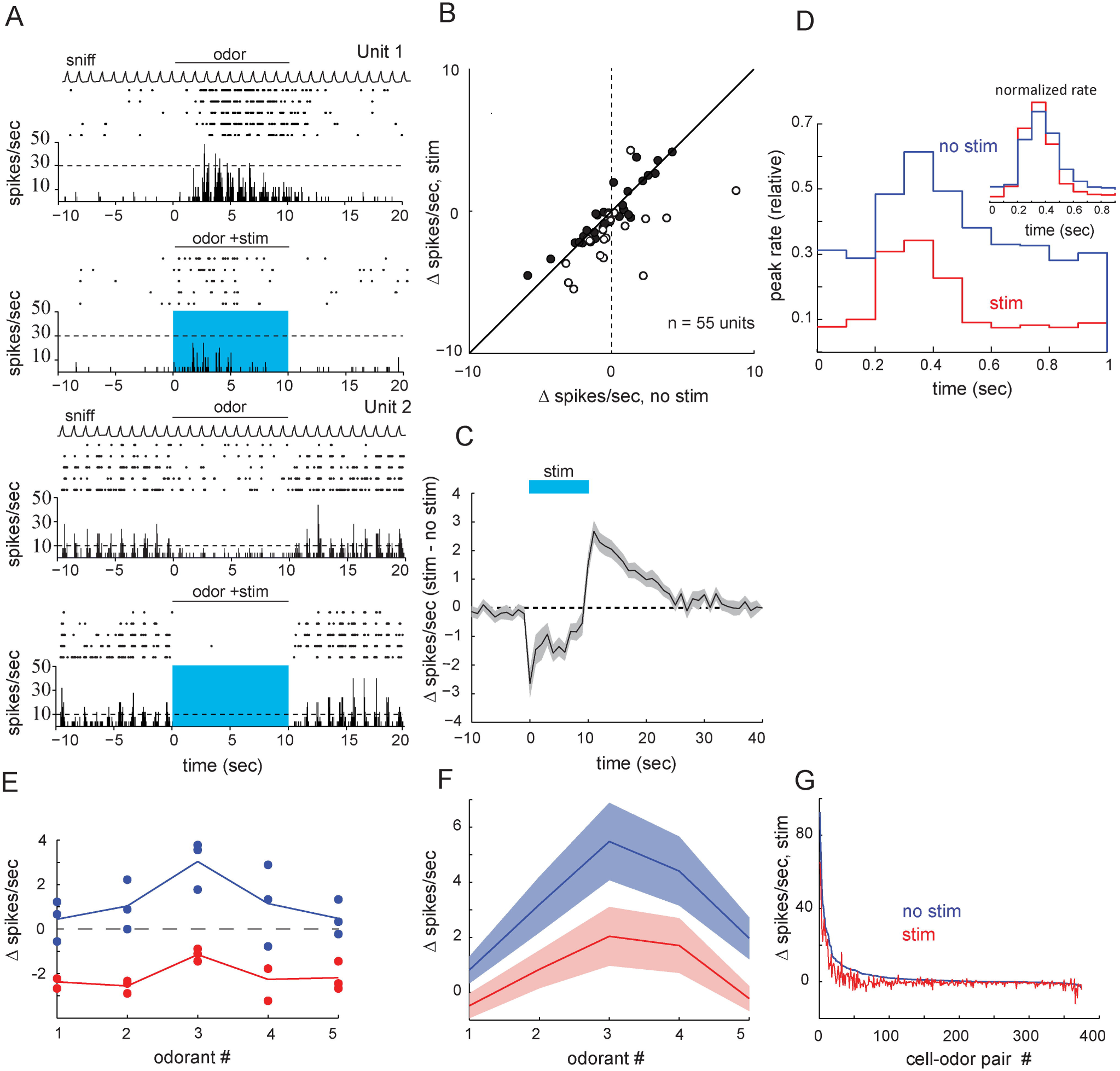
Optogenetic activation of AON axons in the OB inhibits odor-evoked MTC spiking. A. Odorant-evoked MTC spiking is suppressed by optical OB stimulation for neurons that show an excitation (Unit 1,top) or suppression (Unit 2, bottom) of firing rate in response to odorant presentation. B. Plot of odorant-evoked changes in MTC spiking activity (Δ spikes/sniff) in the absence (“no stim”) and during (“stim”) photostimulation (n= 55, 6 mice). Open circles depict units that showed significant light-evoked changes in firing activity when tested on a unit-by-unit basis. 17 of those cells showed a significant decrease, only two showed an increase. C. Time course of the effect of AON fiber activation on odor-evoked MTC spiking, averaged across all units. Shaded area represents the variance (±SEM) around mean. The blue bar shows the time of optical stimulation and simultaneous odorant presentation. Photostimulation leads to a strong reduction of odorant-evoked firing activity. D. Sniff-triggered spike histogram of odorant-evoked MTC spiking during odorant presentation in baseline conditions (“no stim”, blue) and during (“stim”, red) optical OB stimulation. Inset, Sniff-triggered spike histogram normalized to the maximum and minimum bin for the two conditions independently. E. Effect of OB optical stimulation on odorant response spectrum for an example MTC tested with five odorants. Blue, Baseline response; red, response during optical stimulation. Odorants are ordered with the strongest excitatory response in the baseline condition in the middle of the abscissa. The effect of optical stimulation varies with odorant but is always inhibitory. Circles indicate firing rates for each trial; lines connect median responses across all tested trials. F. Effect of OB optical stimulation on odorant response spectrum averaged across all recorded MTCs. Blue, Baseline response; red, response during optical stimulation. Odorants are ordered separately for each unit, with the strongest excitatory response in the baseline condition in the middle of the abscissa. Odorant-evoked spike rates uniformly decreased during light stimulation. Lines connect median responses shaded areas indicates the variance (SEM). G. Odorant response magnitudes (Δ spikes/s) plotted for baseline (blue) and optical stimulation (red) as a function of cell identity, sorted in order of magnitude of excitatory response in baseline conditions. Note that most units show a decrease in odorant-evoked excitation, including those that are suppressed during odorant presentation.

Finally, we tested how AON modulates MTC activity across multiple odorants. Thus, in a separate set of experiments, we measured responses of the same MTC to multiple odorants (see Materials and Methods for odorant panel) with and without optogenetic activation of AON bulbar input (75 units, 3 animal, 375 cell–odor pairs). Odorant-evoked spike rates uniformly decreased during light stimulation for the example unit (Figure 6E) as well as for the average across all recorded units (Figure 6F). 68 % of cell-odor pairs even exhibited an inhibition below pre-odor baseline firing rates. Only 2.6% (10 of the 375 cell-odor pairs), showed an increase (> 1 Δspike/sec) in odorant-evoked spike rates during AON activation (Figure 6G). Thus, the AON strongly inhibits sensory evoked responses across odorants similar to what we observed in the behavioral experiments.

## Discussion

The AON is the most anterior part of olfactory cortex lying directly behind the olfactory bulb and constitutes its largest source of cortical projections. Despite these prominent features, the role of AON in odorant processing as well as the behavioral consequences of its activation have been sparingly investigated. Since the AON itself has been proposed as the first stage of odorant feature convergence, receiving structured sensory information in a bottom up fashion and coding itself for odor objects (Haberly, 2001), we first tested if AON photostimulation can be perceived as an sensory equivalent cue, similar to what has been reported for other sensory processing areas (O’Connor et al., 2013; Guo et al., 2015). In contrast to the predictions from (Haberly, 2001) optogenetic AON stimulation does not elicit a behavioral response in animals trained to report the presence of odorants. Pairing optical stimulation with odor presentation however, reliably suppressed odorant detection of awake freely behaving mice, possibly by modulating odor processing via top-down feedback to the olfactory bulb. The results might point towards the AON as an important modulator of odor perception rather than being a pure odor encoding region. In concordance to the awake data, we could show that activation of AON derived fibers in the OB elicit a strong inhibition of MTC firing on spontaneous as well as on sensory evoked activity making this the first report of matching physiological and behavioral changes in sensory evoked responses using fast, timely controlled optogenetics for the AON so far.

### Optogenetic stimulation of AON

Optogenetic activation was achieved through viral ChR2(H134R)-EYFP expression in the AON of nicotinic receptor alpha 7 (Chrna7)-Cre mice (Rogers et al., 2012b). As shown previously (Rothermel and Wachowiak, 2014) this approach labels a substantial amount of cells within the AON that correspond with morphological descriptions of pyramidal cells (Brunjes and Kenerson, 2010). Expression was observed in all AON areas including pars externa. Recently, it was shown that a subpopulation of AON neurons, AON pars externa, can elicit EPSPs in contralateral OB mitral cells (Grobman et al., 2018). In our experiments we rarely observe excitatory MTC responses in response to AON stimulation. While we cannot rule out that these rare excitatory effects are due to weak AON pars externa stimulation, or reflecting differential AON effects on different OB output types (mitral vs. tufted cells), we argue that the dominant inhibitory response observed in our study is likely mediated by AON pars principalis due to intense viral expression, and the AON fiber placement in this area.

Optogenetic stimulation was performed at the level of the OB for electrophysiogical recordings in order to exclude disynaptic or even polysynaptic effects from other targets of AON projections. In behavior experiments, optogenetic stimulation had to be performed at the level of the AON to exclude insufficient photostimulation, since even monomolecular odorants elicit complex OB activity pattern (Johnson and Leon, 2007; Johnson et al., 2009; Ma et al., 2012; Baker et al., 2019) and stimulation of just the dorsally located AON fibers in the OB would most probably have yielded false negative results. AON fiber stimulation at the level of the OB in the electrophysiological recordings caused clear inhibition on a population level. However, it might seem surprising that this inhibition is not larger given the strong impact of AON stimulation on behavior. This might be partially due to the mode of stimulation (axonal OB stimulation might be less effective compared to somatic AON stimulation), the state-dependent strength of AON input to the OB (AON fibers show a higher activity during wakefulness) (Rothermel and Wachowiak, 2014) as well as different modes of local OB networks (increased activity of inhibitory interneurons during wakefulness (Wachowiak et al., 2013; Cazakoff et al., 2014). Due to the qualitatively matching effects in the two experimental paradigms we hypothesize, that both modes of stimulation elicit the same effects within the olfactory bulb.

### Circuit mechanisms underlying AON modulation

Since AON derived back projections to the OB are glutamatergic in nature, reports of inhibitory AON modulation effects seem counterintuitive. A previous study found a very transient (<5ms) excitation followed by a much stronger and prolonged inhibition in a subpopulation of MTCs (Markopoulos et al., 2012). When adjusting bin sizes to similar values used by (Markopoulos et al., 2012) we also observed this fast excitatory component in our experiments (Figure 5-1). However, Markopoulos et al., 2012 was not able to detect this rapid excitation in odor evoked responses, raising the question of its behavioral relevance.

The circuit mechanisms underlying the strong inhibitory effects by glutamatergic AON fibers on MTCs in the OB remain to be elucidated. Projections from AON to OB are diverse in terms of source and their exact targets in the OB are still not fully known (for review see (Brunjes et al., 2005)). In our experiments, expression was predominately observed in the granule cell and external plexiform OB layers, reaching up to the border of the glomerular layer, consistent with earlier characterizations of AON–OB projections (Reyher et al., 1988). This projection pattern as well, as *in vitro* results from Markopoulos et al., suggest disynaptically mediated inhibition via inhibitory periglomerular neurons in the periglomerular layer and/or granule cells in the granule cell layer.

The AON has been shown to send projections to the ipsilateral as well as to the contralateral OB (Schoenfeld and Macrides, 1984; Shipley and Adamek, 1984; Kay and Brunjes, 2014). In most of our electrophysiological experiments we were unable to distinguish between effects elicited from ipsilateral or contralateral AON projections because of the bilateral injections performed in these animals. However, optogenetically stimulating fibers in the contralateral OB after unilateral AON injection in a subset of electrophysiological experiments, revealed inhibitory effects qualitatively similar to those in bilaterally injected animals. Since contralaterally projecting fibers predominantly target the granule cell layer (Davis and Macrides, 1981; Reyher et al., 1988; Markopoulos et al., 2012; Rothermel and Wachowiak, 2014) these findings as well as the strong inhibition seen in behavioral experiments with unilateral AON stimulation points to granule cells being the major mediators of cortically derived inhibition in the OB, which is in line with findings from previous studies (Boyd et al., 2012; Markopoulos et al., 2012).

### Functional role of modulating AON activity

Our experiments show that the inhibitory influence of AON on olfactory responses can be very strong. Inhibition in the OB has been proposed as a mechanism to sharpen the odor tuning of MTC (Yokoi et al., 1995) however, the observed homogenous suppression during light stimulation is not consistent with an AON mediated sharpening of odor responses. We cannot exclude that optical stimulation is stronger and spatiotemporally more homogenous compared to “intrinsic” AON activity as a result of its input from OB (Lei et al., 2006; Kay et al., 2011), anterior piriform cortex (Haberly and Price, 1978; Luskin and Price, 1983; Haberly, 2001; Hagiwara et al., 2012), amygdala (De Carlos et al., 1989; Gomez and Newman, 1992; Canteras et al., 1995; Petrovich et al., 1996), basal forebrain (Broadwell and Jacobowitz, 1976; Luiten et al., 1987; De Carlos et al., 1989; Carnes et al., 1990; Gaykema et al., 1990; Zaborszky et al., 2012; Gielow and Zaborszky, 2017) hippocampus, (Swanson and Cowan, 1977; van Groen and Wyss, 1990; Aqrabawi and Kim, 2018b) and medial prefrontal cortex (Sesack et al., 1989). While speculating about “intrinsic” modes of AON activation is complicated by these extensive connection with olfactory as well as non-olfactory centers and therefore out of the scope of this study, we could show that photostimulation did not affect general performance, as ChR2 mice were able to perform non olfactory tasks during laser stimulation (data not shown). Since odor specific “feedback” activation pattern on AON top-down fibers in the OB have been demonstrated (Rothermel and Wachowiak, 2014) one can speculate that the AON might be supporting fast adaptation or adjusting the dynamic range of MTCs to strong odorants. Additionally, there might also be “state-dependent” AON activation by higher brain areas which might be spatiotemporally much broader, thereby potentially more closely reflecting the here applied optical stimulation. In line with our results, activation of a subregion of the AON (medial AON, potentially driven by hippocampal inputs) using a chemogenetic approach was recently reported to reduce olfactory sensitivity as well as to impair the performance of olfaction-dependent behaviors (Aqrabawi et al., 2016). However, due to the slow nature of the chemogenetic approach, the temporal component of this effect could not be investigated.

Overall our results clearly demonstrate the strong modulatory potential of the AON on odor processing. We show that AON stimulation in awake mice suppresses odor responses irrespective of odor identity or odor concentration. Furthermore AON derived fiber activation within the OB elicited a strong inhibition of spontaneous as well as odor evoked MTC firing with even strongly odorant activated units being inhibited below pre odor baseline firing rate. Together this builds the first report of matching physiological and behavioral changes in sensory evoked responses using fast, timely controlled optogenetics for the AON and implicates the AON as a potential gatekeeper of olfactory information.

## Acknowledgments

We thank Scott W. Rogers and Petr Tvrdik for kindly providing the Chrna7-Cre mice. Chrna7-Cre mice were generated with funding from NIH AG017517 to S. Rogers. We thank the technical workshop at the institute for biology II, RWTH Aachen for excellent technical support. We thank Matt Wachowiak, Jeremy C. McIntyre and all members of the M.R. lab members for helpful discussion and comments on the manuscript. Initial data were acquired in Utah. We thank Drs. Looger, Akerboom and Kim and the Genetically Encoded Calcium Indicator (GECI) Project at Janelia Farm Research Campus in collaboration with Penn Vector Core for providing with GCaMP-expressing viruses. This work was supported by funding from DFG (RO4046/2-1 and /2-2, Emmy Noether Program [to MR] and the Research Training Group 2416 “MultiSenses – MultiScales: Novel approaches to decipher neural processing in multisensory integration”

